# A comparative study of SVDquartets and other coalescent-based species tree estimation methods

**DOI:** 10.1101/022855

**Authors:** Jed Chou, Ashu Gupta, Shashank Yaduvanshi, Ruth Davidson, Mike Nute, Siavash Mirarab, Tandy Warnow

**Affiliations:** Department of Mathematics, University of Illinois Urbana-Champaign, 1409 W. Green Street, 61801 Urbana, IL, USA; Department of Computer Science, University of Illinois Urbana-Champaign, 201 N Goodwin Ave, 61801 Urbana, IL, USA; Department of Statistics, University of Illinois Urbana-Champaign, 725 South Wright Street, 61820 Champaign, IL, USA; Department of Electrical and Computer Engineering, University of California San Diego, 9500 Gilman Drive, 92093 La Jolla, CA, USA; Department of Computer Science, University of Texas at Austin, 2317 Speedway, Stop D9500, 78712 Austin, TX, USA

**Author notes:** Correspondence Department of Computer Science, University of Illinois Urbana-Champaign, 201 N Goodwin Ave, 61801 Urbana, IL, USA.

**Keywords:** species tree inference, SNP, multilocus, SVDquartets, QMC, ASTRAL, NJst, RAxML

## Abstract

**Background:** Species tree estimation is challenging in the presence of incomplete lineage sorting (ILS), which can make gene trees different from the species tree. Because ILS is expected to occur and the standard concatenation approach can return incorrect trees with high support in the presence of ILS, “coalescent-based” summary methods (which first estimate gene trees and then combine gene trees into a species tree) have been developed that have theoretical guarantees of robustness to arbitrarily high amounts of ILS. Some studies have suggested that summary methods should only be used on “c-genes” (i.e., recombination-free loci) that can be extremely short (sometimes fewer than 100 sites). However, gene trees estimated on short alignments can have high estimation error, and summary methods tend to have high error on short c-genes. To address this problem, Chifman and Kubatko introduced SVDquartets, a new coalescent-based method. SVDquartets takes multi-locus unlinked single-site data, infers the quartet trees for all subsets of four species, and then combines the set of quartet trees into a species tree using a quartet amalgamation heuristic. Yet, the relative accuracy of SVDquartets to leading coalescent-based methods has not been assessed.

**Results:** We compared SVDquartets to two leading coalescent-based methods (ASTRAL-2 and NJst), and to concatenation using maximum likelihood. We used a collection of simulated datasets, varying ILS levels, numbers of taxa, and number of sites per locus. Although SVDquartets was sometimes more accurate than ASTRAL-2 and NJst, most often the best results were obtained using ASTRAL-2, even on the shortest gene sequence alignments we explored (with only 10 sites per locus). Finally, concatenation was the most accurate of all methods under low ILS conditions.

**Conclusions:** ASTRAL-2 generally had the best accuracy under higher ILS conditions, and concatenation had the best accuracy under the lowest ILS conditions. However, SVDquartets was competitive with the best methods under conditions with low ILS and small numbers of sites per locus. The good performance under many conditions of ASTRAL-2 in comparison to SVDquartets is surprising given the known vulnerability of ASTRAL-2 and similar methods to short gene sequences.

## Background

Estimating a species tree from multi-locus sequence data is complicated by biological processes such as gene duplication and loss, hybridization, and incomplete lineage sorting, which make true gene trees different from the overall true species tree. Incomplete lineage sorting (ILS) [1], where gene lineages from two taxa fail to coalesce in the most recent ancestor, is one of the common sources of discordance between gene trees and species trees [2] and is statistically modeled by the multi-species coalescent [3].

Methods for estimating species trees in the presence of ILS have been developed that are provably statistically consistent under the multi-species coalescent model, which means that they will converge in probability to the true species tree as the number of loci and sites per locus increase [4]. The most popular such techniques are summary methods, in which an alignment is estimated on each locus, a gene tree is estimated on each alignment, and then the resulting gene trees are combined into a species tree. Examples of summary methods include ASTRAL [5], ASTRAL-2 [6], MP-EST [7], the population tree from BUCKy [8], and NJst [9]. Statistically consistent co-estimation of gene trees and species trees is possible [10, 11], but these methods are much more computationally intensive than the most popular summary methods and so are not used on large-scale phylogenomic datasets [12, 13, 14].

The most common approach for estimating species trees is concatenated analysis using maximum likelihood (CA-ML), in which alignments for each locus are aggregated into a supermatrix and a species tree is estimated using a maximum likelihood (ML) method such as RAxML [15] or FastTree-2 [16], under a statistical model in which all sites evolve identically and independently (*i.i.d.*) down a single model tree. However, CA-ML is not statistically consistent under the multi-species coalescent and can converge to a tree other than the species tree (i.e., be positively misleading) [17].

Because concatenation can be positively misleading, the interest in using coalescent-based species tree estimation methods has increased. Since summary methods are able to analyze large datasets (some are sufficiently fast that they can analyze datasets with 1000 species and 1000 genes [6]), most coalescent-based analyses of biological datasets have been performed using summary methods [18, 19]. Several summary methods have good accuracy on small datasets (with up to 10 taxa), including ASTRAL [5], ASTRAL-2 [6], NJst [9], BUCKy-pop [8, 20], and MP-EST [7]. ASTRAL-2 [6] and NJst [9] generally dominate MP-EST on larger datasets in terms of accuracy, and BUCKy-pop and MP-EST are both too slow to run on datasets with many taxa. More generally, only ASTRAL-2 and NJst are fast enough to run on very large datasets with high accuracy [6, 21, 20].

Since the multi-species coalescent model requires representing each gene by a single tree, it allows no recombination events inside a gene [22], and so the guarantees of statistical consistency can fail in the presence of recombination. For this reason, some practitioners have argued [23] that only recombination-free alignments (i.e., coalescent-genes, or “c-genes”) should be used with summary methods. However, c-genes can be very short (less than 100 sites), and depending on the scope of the taxonomic study are likely to be closer to a single site than 100 [23]. Because summary methods are sensitive to gene tree estimation error [24, 25, 26, 14], which is more likely to occur on short alignments, the utility of summary methods for phylogenomic species tree estimation is questioned due to this perceived need to constrain the sequence length for every locus [23].

The relative performance of concatenation and summary methods is clearly impacted by the amount of ILS, so that concatenated analyses are often more accurate than coalescent-based methods if the ILS level is low enough, and less accurate for high levels of ILS [26, 14]. However, the relative performance is also impacted by gene alignment lengths (shorter gene sequence alignments tend to produce higher gene tree estimation error, and hence higher species tree estimation error for summary methods) [24, 14, 27]. Finally, the number of genes and taxa also have an impact on the relative performance [14, 27]. Thus, despite the theoretical advantages of coalescent-based summary methods, there is a heated debate about whether summary methods or a maximum likelihood concatenated analysis would more accurately estimate phylogenies under biologically realistic conditions [23, 24, 2, 28, 29].

An alternative approach for coalescent-based species tree estimation has been proposed that avoids estimating trees on each locus, and hence bypasses the issue of gene tree estimation error. These methods, which we refer to as “single-site” methods, examine the single-site patterns, and use these patterns to estimate the species tree in a statistically consistent way. The first such method, SNAPP [30], required biallelic (two-state) data from unlinked single-nucleotide polymorphisms (SNPs), and employed a Bayesian MCMC algorithm. More recently, Chifman and Kubatko introduced SVDquartets, a single-site method that can handle nucleotide data. Using algebraic statistics and singular value decomposition, Chifman and Kubatko proved that under the multi-species coalescent and with the assumption of a strict molecular clock (i.e., that the rate of sequence evolution per unit time is constant throughout the model gene tree), an unrooted species tree on four taxa is generically identifiable from site pattern probabilities at the leaves of the tree [31]. The SVDquartets algorithm takes unlinked multi-locus data for a set of four taxa as input and assigns a score to each of the three possible quartet topologies. The quartet topology with the lowest “SVD score” is selected as the true topology for that quartet [32].

Since SVDquartets just computes quartet trees, a quartet tree agglomeration technique is needed to combine the quartet trees on every four species into a species tree on the full set of taxa. Chifman and Kubatko [31] suggested the use of Quartet Max-Cut (QMC) [33], a heuristic for the NP-hard Max Quartet Compatibility problem [34, 35]. However, SVDquartets has also been implemented in PAUP* [36], which uses a variant of Quartet FM [37] to combine quartet trees into a species tree, and is the implementation currently recommended by the developers of SVDquartets.

To our knowledge, the accuracy of species tree estimation using SVDquartets followed by any quartet amalgamation method has not yet been explored in comparison to summary methods or concatenation. In this study, we address the following questions:

1. How does SVDquartets+PAUP* compare to ASTRAL-2 and NJst, two of the best performing statistically consistent summary methods?
2. How do the statistically consistent methods we study compare to a concatenated analysis using maximum likelihood?
3. How do all the methods perform on short sequences?

We ran ASTRAL-2 and NJst on gene trees estimated by FastTree-2 [16], a maximum likelihood method for species tree inference with similar accuracy to RAxML [38]. We also ran an unpartitioned concatenated maximum likelihood analysis using RAxML. We omit MP-EST from the evaluation on simulated data because ASTRAL-2 generally dominates MP-EST in terms of accuracy and running time [6, 26, 27]. As all species trees we estimated are fully resolved (i.e., bifurcating), we evaluate species tree estimation methods using the Robinson-Foulds (RF) [39] error rate, also known as the normalized bipartition distance.

We used previously studied simulated datasets and simulated new datasets as well to evaluate the performance of these methods. Table 1 presents a summary of these datasets, which vary in number of taxa (11 to 37), ILS level (reflected in the average topological distance between true gene trees and true species tree) and whether sequence evolution is under a strict molecular clock (which SVDquartets assumes). See Methods for additional details.

**Table 1.** Empirical statistics of simulated datasets. We show ILS level (given in terms of average distance between true gene trees and true species trees), number of sites per gene, number of genes, whether a strict molecular clock applies, and number of replicate species trees. The number of taxa is indicated by the dataset name.

| Summary of Simulated Datasets |  |  |  |  |  |  |
| --- | --- | --- | --- | --- | --- | --- |
| Dataset | ILS level (AD) | # Sites | # Genes | Clock | # Reps. | Ref. |
| 11-taxon M1 | 15.5% | 10-200 | 100-1000 | No | 50 | (new) |
| 11-taxon M2 | 38.3% | 10-200 | 100-1000 | No | 50 | [27] |
| 11-taxon M3 | 66.3% | 10-200 | 100-1000 | No | 50 | (new) |
| 11-taxon M4 | 85.0% | 10-200 | 100-1000 | No | 50 | [27] |
| 15-taxon | 82% | 10-200 | 100-1000 | Yes | 10 | [27] |
| 37-taxon Mammalian | 18% | 10-200 | 50-200 | No | 20 | [26] |

Previous studies have only examined coalescent-based methods on gene sequence alignments with at least 100 sites. To understand performance on short sequences, we shortened each of the gene alignments by sampling the first 10, 25, 50, 100, or 200 sites from the gene sequence alignments.

In addition, we ran SVDquartets+PAUP* on a well-studied mammalian biological dataset from [40] (after removing erroneous genes) with 37 taxa, and compared the output species tree to previously published [14, 27] trees computed using ASTRAL-2, MP-EST, and concatenation with maximum likelihood.

## Results

### Results on 11-taxon datasets

The results on the 11-taxon datasets are shown in Figure 1, varying the ILS level from low (model M1) to very high (model M4). For all methods, tree estimation error rates reduce as the number of genes or sites per gene increase, while they increase as the ILS level increases.

**Figure 1.**
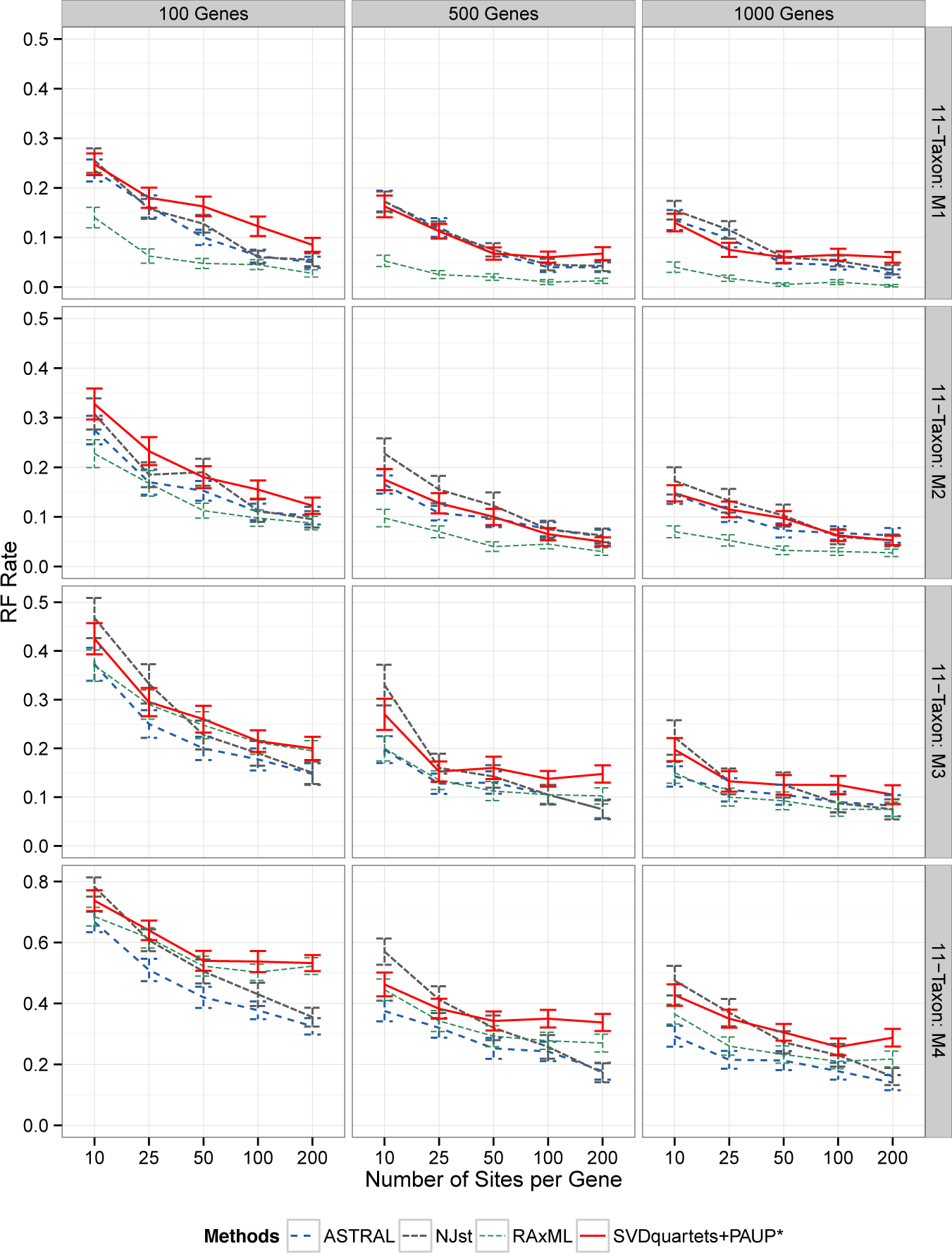
Results on the 11-taxon simulated datasets. We show mean RF rates with standard error bars for 50 replicates using all four methods (RAxML shows concatenation). The rows are for four 11-taxon (10 ingroup taxa and one outgroup taxon) model conditions with varying ILS levels, ranging from very low (M1) to very high (M4). The columns are for the different numbers of genes. Within a subfigure, we show results with changing numbers of sites per locus (10-200). Note that the y-axis range changes for the fourth row, due to the much higher error rates under the highest ILS model condition. Sequence evolution on these datasets deviates from the strict molecular clock.

However, the relative performance between methods depends on the model condition. For example, CA-ML (in purple) is the most accurate for models M1 and M2 for all numbers of genes and numbers of sites per gene. On model M3, CA-ML is one of the least accurate methods for the 100-gene datasets, but close to the most accurate on the datasets with 500 or 1000 genes. On model M4, with the highest ILS, CA-ML is among the least accurate methods on the 100-gene datasets, but intermediate on the 500- and 1000-gene datasets. Thus, CA-ML has excellent accuracy on the two lower ILS model conditions and then average accuracy on the two higher ILS model conditions.

The remainder of this discussion focuses on a comparison of the coalescent-based methods (i.e., ASTRAL-2, NJst, and SVDquartets+PAUP*). Since the relative performance is impacted by the ILS level, we discuss each model in turn, beginning with model M1 (lowest ILS).

On model M1, differences between methods were small, and in general all methods had very good accuracy. On the datasets with 10 or 25 sites per gene, all methods had nearly identical accuracy. SVDquartets+PAUP* was slightly more accurate than ASTRAL-2 and NJst with 500 10-site genes and 1000 10-site or 25-site genes. However, on datasets with 50 to 200 sites per gene, SVDquartets+PAUP* was sometimes less accurate than the other methods.

Results on the models with higher ILS show that methods varied in their robustness to ILS, so that ASTRAL-2 had the greatest robustness to ILS, and there were bigger differences between methods in the presence of high ILS. On the datasets with only 10 sites per gene, ASTRAL-2 had the best accuracy, followed by SVDquartets+PAUP*, and then by NJst. However, SVDquartets+PAUP* had the least accurate results of all methods on these higher ILS models, especially when the number of sites per gene was 50 or greater. Overall, ASTRAL-2 has better accuracy than SVDquartets+PAUP*, and as the ILS increases, the gap between ASTRAL-2 and SVDquartets+PAUP* also increases.

We evaluated the statistical significance of the difference in mean RF rates between SVDquartets+PAUP* and ASTRAL-2 at each level of ILS, and we further test the hypothesis that the interaction between level of ILS and method is non-zero (that is, that the difference in mean RF rates changes as ILS increases). For this we use an ANOVA test with a linear model, where the level of ILS, number of genes and number of sites per gene are all treated as (ordinal) categorical variables. For the former test, we conduct the test simultaneously for the 11-taxon datasets at all four levels of ILS (M1-M4), and for the latter at the three levels (M2-M4) that represent a change in ILS. Under this procedure, we reject the null hypothesis that the two methods have equivalent mean RF rates at any of the four levels of ILS (Table 2). We further reject the null hypothesis that the methods degrade in performance at the same rate from M2 to M3 and from M3 to M4.

**Table 2.** *p*-values for statistical tests. We use ANOVA to compare mean tree error of ASTRAL-2 and SVDquartets+PAUP* on different model conditions (left column) and also to test whether the differences are impacted by changing levels of ILS (right column). All seven p-values are corrected for multiple hypothesis testing using a Benjemani-Hochberg procedure [41] (*n* = 7). Italics indicate significance (*α* =.05).

| Mean RF Rate |  | Interaction |  |
| --- | --- | --- | --- |
| ILS | *p* | $\Delta$ ILS | *p* |
| M1 | <.0001 | M1-M2 | 0.946 |
| M2 | <.0001 | M2-M3 | 0.0403 |
| M3 | 0.0403 | M3-M4 | <0.0001 |
| M4 | 0.0403 |  |  |

### Results on the 15-taxon datasets

The 15-taxon datasets are the only datasets we explored where sequences evolve under a strict molecular clock. As with the 11-taxon model conditions, errors reduced with increasing numbers of sites per gene or numbers of genes (Fig. 2). CA-ML and SVDquartets+PAUP* nearly always had higher error rates than ASTRAL-2 and NJst for all numbers of genes and sites per gene (the only exception to this is on 100 genes with 10 sites per gene, where NJst and CA-ML had the same error). The relative accuracy of ASTRAL-2 and NJst depended on the specific model condition (number of genes and number of sites), and neither outperformed the other.

**Figure 2.**
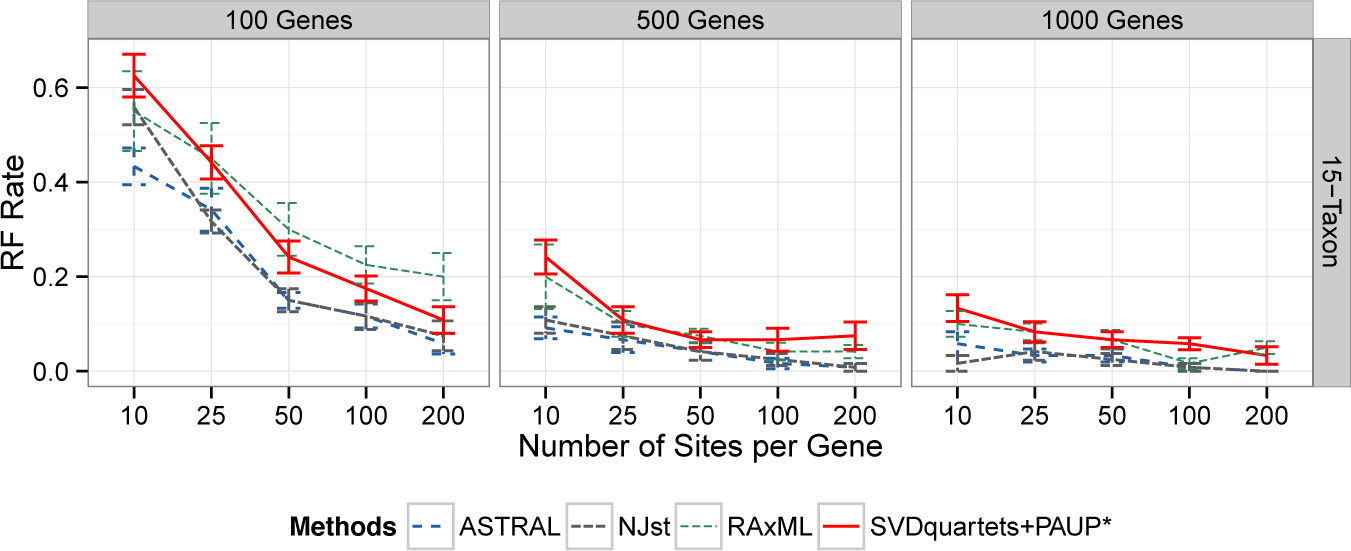
Results on the 15-taxon simulated datasets. We show mean RF error rates with standard error bars over 10 replicates, for 100 to 1000 genes. Within a subfigure, we show results with changing numbers of sites per locus (10-200). The 15-taxon model tree is a caterpillar (pectinate tree) with very short internal branches, and these datasets have very high ILS. Sequence evolution on these datasets is under the strict molecular clock.

**Figure 3.**
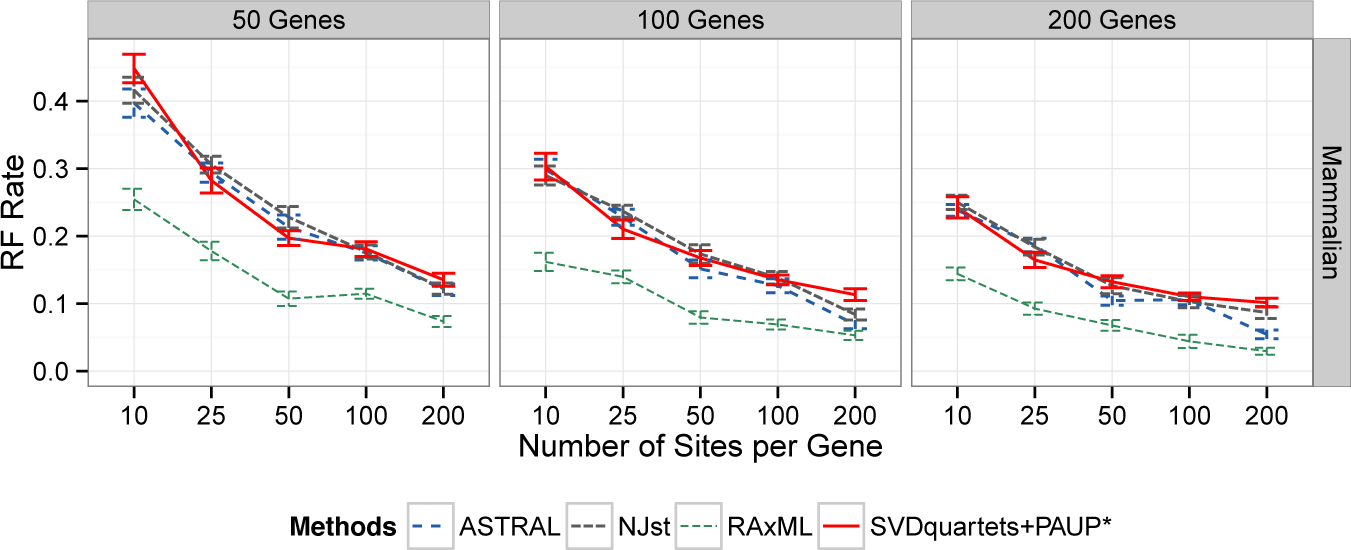
Results on the low ILS mammalian simulated datasets. We show mean RF error rates with standard error bars over 20 replicates, for 50 to 200 genes. Within a subfigure, we show results with changing numbers of sites per locus (10-200). The mammalian simulation is for 37 species, and is based on an MP-EST analysis of a biological dataset with reduced ILS (produced by doubling the species tree branch lengths). Sequence evolution in this simulation deviates from the strict molecular clock.

### Results on the mammalian simulated datasets

We explored performance on a simulated dataset based on a prior MP-EST analysis of a 37-taxon mammalian biological dataset studied in [40] with 50 to 200 genes and a low rate of ILS. This dataset has reduced ILS relative to the original biological dataset from [40], and so represents a relatively easy model condition. Results on these data (Fig. 3) show that error rates decreased for all methods as the number of sites per locus or number of loci increased (as observed in the other model conditions). CA-ML was substantially more accurate than the coalescent-based methods, with the largest differences on the datasets with small amounts of data.

On these mammalian simulated data, concatenation using unpartitioned maximum likelihood was by far the most accurate method, with big differences between concatenation and the coalescent methods for small amounts of data. The differences between the coalescent-based methods were generally small and depended on the number of sites per locus and number of loci, but the most accurate method was always either ASTRAL-2 or SVDquartets+PAUP*. On 10 sites, all the methods had very close accuracy, with a slight advantage for ASTRAL-2 on the 50-gene condition. However, on 25 sites, SVDquartets+PAUP* was the most accurate method (even though differences were small). For larger numbers of sites per gene, ASTRAL-2 was the most accurate method.

#### SVDquartets+PAUP* on the mammalian biological dataset

The mammalian biological dataset has been analyzed in prior studies, with trees computed using CA-ML and also MP-EST and ASTRAL-2, each with or with-out statistical binning (both weighted and unweighted) [14, 27]. Statistical binning (weighted and unweighted) are new techniques aimed at improving the estimation of gene trees in a multi-locus phylogenomic analysis, when individual genes have low signal, and have both been shown to improve the accuracy of summary methods [27]. Here we examine the differences between the tree obtained by SVDquartets+PAUP* and these previously published trees.

The species tree obtained by SVDquartets+PAUP* has very high bootstrap support on most branches, but has one branch with very low support (only 35%); see Figure 4. The branch with very low support should be collapsed, leaving a soft polytomy between Cetartiodactyla, Chiroptera, and ((Felis catus, Canis familiaris), Equus caballus).

**Figure 4.**
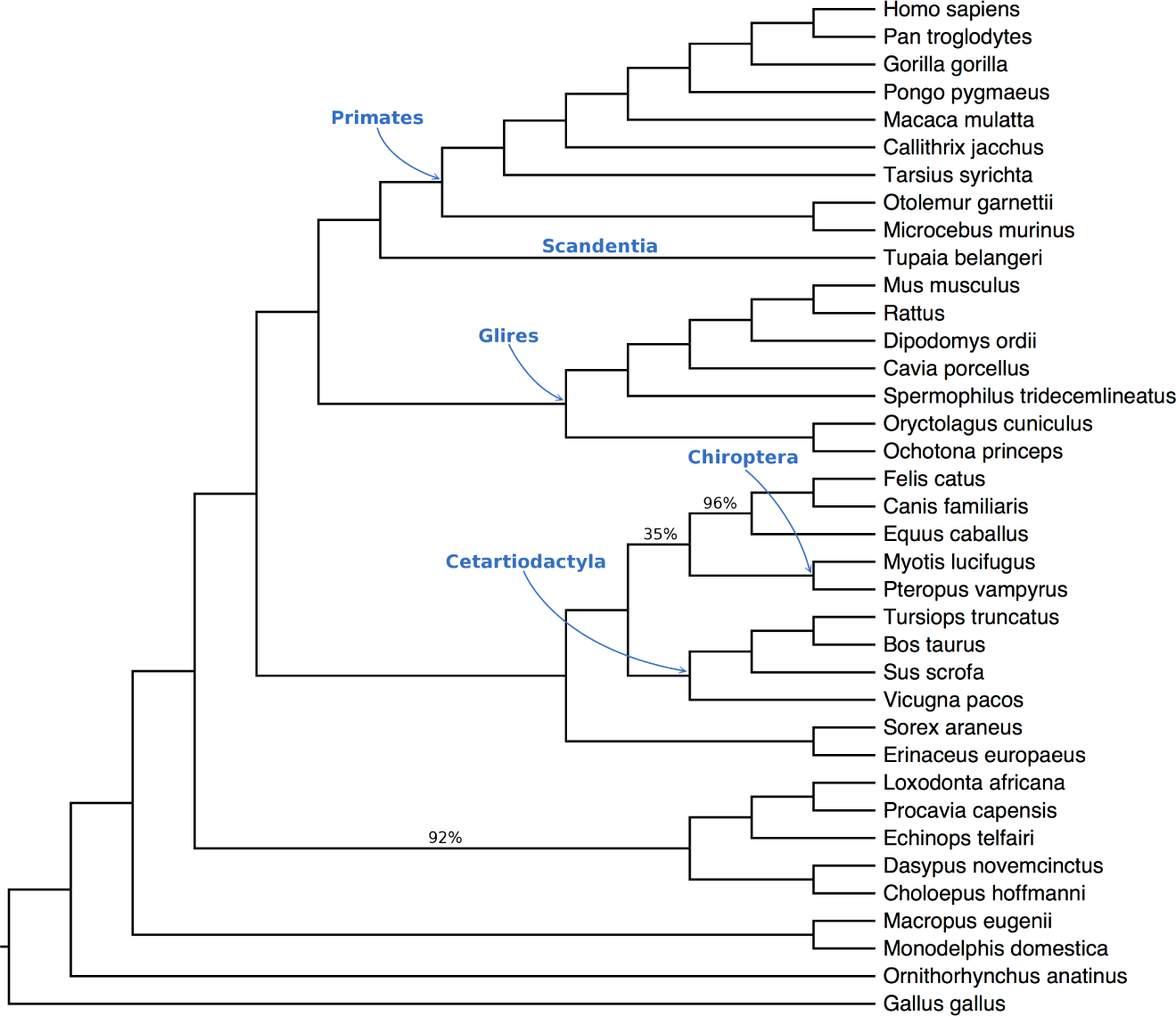
The SVDquartets+PAUP* tree on the 37-taxon 424-gene mammalian dataset from Song et al. [40]. This tree has one branch with very low support, and so does not resolve the relationship between Cetartiodactyla, Chiroptera, and the clade ((Felis catus, Canis familiaris), Equus caballus). Labels on branches indicate bootstrap support, but support values of 100% are not shown.

The only two topological differences between the species trees output by the various methods concern the placement of Scandentia (represented by Tupaia belangeri), and the topology of the clade involving Cetartiodactyla, Chiroptera, and the clade ((Felis catus, Canis familiaris), Equus caballus). ASTRAL-2 (with weighted and unweighted binning, and without binning), MP-EST (with weighted and un-weighted binning), and CA-ML all placed Scandentia sister to Glires. In contrast, SVDquartets+PAUP* and unbinned MP-EST placed Scandentia sister to Primates. The placement of Scandentia is debated, and so it is not clear which of these relationships is correct [27, 42, 43].

All three possible topologies for Cetartiodactyla, Chiroptera, and the clade ((Felis catus, Canis familiaris), Equus caballus) were obtained by the various methods. The three ways of running ASTRAL-2 and MP-EST (with weighted and unweighted binning, and without binning) all resolved the populations Cetartiodactyla, Chiroptera, and ((Felis catus, Canis familiaris), Equus caballus) by placing Cetartiodactyla as siblings with ((Felis catus, Canis familiaris), Equus caballus), each with support of around 80%. In contrast, CA-ML placed Cetartiodactyla sister to Chiroptera with 76% support [14]. Finally, as noted, although SVDquartets+PAUP* placed Chiroptera sister to ((Felis catus, Canis familiaris), Equus caballus), the branch leading to this pair had an extremely low bootstrap support of 35%, which is best considered as not resolving the relationship between Cetartiodactyla, Chiroptera, and ((Felis catus, Canis familiaris), Equus caballus).

## Discussion

To the best of our knowledge, this study is the first to compare species tree estimation methods on short gene sequences (with fewer than 100 sites), and the first to explore SVDquartets on simulated data in comparison to other coalescent-based methods and to concatenation. Many of the trends we noted have been observed in other studies. For example, species tree estimation error tends to decrease for all methods with decreases in the ILS level, increases in gene sequence lengths, and increases in numbers of genes. This study also confirmed that using unpartitioned CA-ML is often more accurate than coalescent-based methods when the ILS level is low enough, but that high ILS levels reverses this relationship. As seen in [6], ASTRAL-2 is typically (but not always) more accurate than NJst, and NJst tends to have reduced accuracy under conditions with high ILS compared to ASTRAL-2. While these trends are not novel [14, 27, 23, 14], this study confirms these results on new datasets, and hence provide additional support for these findings.

In general, ASTRAL-2 tended to be more accurate than SVDquartets+PAUP*, but there were exceptions that could be characterized by small numbers of sites, large numbers of genes, and ILS levels that were not too high. In addition, the results on the 11-taxon datasets suggest that ILS level impacts SVDquartets+PAUP* more significantly than it does ASTRAL-2, so that SVDquartets+PAUP* can be more accurate than ASTRAL-2 on low ILS model conditions and then less accurate than ASTRAL-2 on high ILS conditions. SVDquartets+PAUP* was also much less accurate than ASTRAL-2 on the 15-taxon datasets, which have very high ILS, adding additional support to the hypothesis that SVDquartets+PAUP* degrades on high ILS conditions.

From a purely theoretical perspective, SVDquartets assumes that sequence evolution obeys the strict molecular clock, but this study shows it has fairly good accuracy under the model conditions that deviate from this assumption. Hence, in practice, SVDquartets may be robust to violations of the molecular clock hypothesis.

While SVDquartets+PAUP* had good accuracy on these data, it did not tend to have better accuracy than the competing methods, except as discussed above. However, SVDquartets+PAUP* is a new type of approach, and the design space has not been fully explored. Therefore, it is not clear if the cases where SVDquartets+PAUP* is less accurate than ASTRAL-2 are due to the way that quartet trees are computed by SVDquartets, or by how PAUP* agglomerates them into a species tree. However, the good performance of SVDquartets+PAUP* on conditions of low to moderate ILS suggests that the quartet amalgamation technique in PAUP* may be highly effective when there is not too much gene tree discord.

An important observation across our study was that with very short sequences, all methods had very high error rates, and that SVDquartets+PAUP* did not have an advantage over the best summary methods under these conditions. Thus, an attempt to avoid recombination comes at the cost of reduced accuracy for all methods. Whether it is necessary or not to restrict the data to c-genes, however, is debated, as some studies have shown that summary methods are robust to the presence of recombination in gene trees [44]. In addition, the accuracy of the naive binning approach in experimental studies [45] is compatible with those conclusions. An alternative to choosing very small “c-genes” is to use longer genes, hoping that effects of recombination are minimal. However, more research is needed to better understand the extent of robustness of summary methods to recombination events.

## Conclusions

This study was motivated by the challenge of estimating species trees in the presence of gene tree heterogeneity due to incomplete lineage sorting (ILS) [46, 24]. Although some very sophisticated Bayesian species tree estimation methods, such as *BEAST [11], have been developed, for computational reasons only summary methods (which estimate species trees by combining estimated gene trees) have become widely used in phylogenomics. Yet, there is a large controversy around the use of summary methods, centering on the observation that recombination-free sequence alignments can be very short, and that standard summary methods can produce species trees with reduced accuracy when the gene trees have reduced accuracy resulting from insufficient sequence length [45, 14, 25]. Furthermore, it is not known whether the standard summary methods are even statistically consistent when the genes have bounded sequence lengths [24]. Hence, some [48] have suggested that summary methods should not be used unless they are established to have better accuracy than alternative methods (such as concatenation, which is not even statistically consistent in the presence of ILS) under biologically realistic conditions (i.e., either sequences that are short enough to be recombination-free, or on datasets in which the gene sequences evolve with recombination).

The SVDquartets method was developed to address this challenge. Instead of using estimated gene trees, it estimates the species tree directly from the site patterns, and hence bypasses the impact of gene tree estimation error. Thus, SVDquartets+PAUP* (the implementation of SVDquartets within PAUP*) is a species tree estimation method that could, conceivably, provide much improved species tree estimation accuracy compared to standard summary methods and concatenation.

While the study we have presented was limited in scope, some trends clearly emerge. First, the relative performance of SVDquartets+PAUP*, ASTRAL-2, and CA-ML depend on the amount of ILS and sequence length per locus, so that SVDquartets+PAUP* can be slightly more accurate than ASTRAL-2 under conditions where both ILS levels are low and sequences are short. However, in most conditions, such as when the level of ILS was high, or when many sites were available, the best summary methods tended to outperform SVDquartets+PAUP*. The comparison to concatenation is also interesting: concatenation using an unpartitioned maximum likelihood analysis is not statistically consistent in the presence of ILS, but it seems to have very good accuracy under low ILS model conditions. SVDquartets+PAUP* is not generally as accurate as concatenation under the low ILS model conditions, but can be more accurate under the higher ILS model conditions.

This study explored the performance of SVDquartets+PAUP* in comparison to other coalescent-based methods and to concatenation. A better understanding of the relative accuracy of methods will require a wider range of methods, including fully partitioned maximum likelihood (which has different statistical properties from unpartitioned maximum likelihood) [4]. It would also be interesting to evaluate coalescent-based methods that co-estimate gene trees and species trees, such as *BEAST, as well as summary methods that use gene trees computed using Bayesian methods, such as PhyloBayes-3 [49].

The statistical guarantees for SVDquartets requires a strict molecular clock, a property that is not likely to hold on most biological datasets, especially when the species are not very closely related, and when loci are sampled from throughout the genomes. For this reason, our study focused on datasets that violate the strict molecular clock. However, the accuracy of SVDquartets could improve under a strict molecular clock, and future work should also investigate this possibility.

Finally, the results shown here focused on the implementation of SVDquartets within PAUP*, and improved empirical performance might be obtained through the development of new heuristics for the optimization problem. In addition, since the PAUP* heuristic is not guaranteed to find an optimal solution and is not even guaranteed to find the compatibility tree when all the quartet trees produced by SVDquartets are identical to the species tree, it is not clear whether SVDquartets+PAUP* is statistically consistent under the multi-species coalescent model. Thus, improved theoretical guarantees might also result through the use of alternative quartet amalgamation methods.

## Methods

### Datasets

We explored results on a collection of datasets, including one 37-taxon biological dataset and several simulated datasets (see Table 1 for a description of the simulated datasets). The ILS level of a dataset (denoted by AD) was measured by the average topological distance between the model gene trees and the model species tree, where the distance between two trees is the number of bipartitions unique to the two trees divided by 2(*n* – 3), where *n* is the number of taxa, expressed as a percentage. This distance is also known as the Robinson-Foulds rate, expressed as a percentage.

#### Mammalian biological dataset

This 37-taxon dataset of mammals with 447 gene alignments was studied by Song et al. [40]. As noted in [14], 23 of the 447 loci were removed because 21 contained mislabeled sequences and two were outliers. We ran SVDquartets+PAUP* with 100 bootstrap replicates on the 424 remaining gene alignments. This produced a set of 100 bootstrap replicate species trees and their greedy consensus tree.

#### 11-taxon dataset

The 11-taxon datasets were partially studied in [27], but we added two new model conditions to this dataset (M1 and M3). In total, our collection has 200 replicate species trees generated under four model conditions (M1, M2, M3, and M4). These data were simulated with SimPhy [50] using parameters given in [27] and scripts that are available in our github repository. The species trees were simulated using the Yule process, with the birth rate set to 0.000001 per generation; hence, the model species trees for these 11-taxon datasets range in topology, but are neither perfectly balanced nor pectinate (the two extremes of tree shape). The four model conditions all had a population size of 400k, but differed in terms of their tree length (5400k, 1800k, 600k, and 200k respectively for M1-4). The change in tree length results in change in ILS levels between M1 and M4, with AD ranging from very low (15.5% for M1) to very high (85.0% for M4).

#### 15-taxon dataset

This dataset was studied in [27]. The simulated species tree is pectinate (a tree of the form (*s*1, (*s*2, (*s*3, (*s*4, *…*))))) on 15 leaves with very short internal branches. These sequences evolve under a strict molecular clock, and with very high ILS (AD=82%).

#### Mammalian simulated dataset

The mammalian simulation is based on a model species tree previously computed using MP-EST on the 37-taxon mammalian biological dataset from [40]. This simulated dataset has been studied in [14, 26] and corresponds to the 2X condition, where all model tree branches are multiplied by two to reduce ILS. The average distance between true gene trees and the model species tree was moderate, at AD=18%.

#### Shortening alignments and processing data

Each of the original datasets we obtained had sequences that varied in length, but all were at least 200 sites. Given a set of gene alignments, we kept the first *K* sites from each alignment, allowing *K* to vary between 10, 25, 50, 100, and 200. Thus, for a given alignment, the corresponding shortened alignment of length 10 is contained in the shortened alignment of length 25, which is contained in the shortened alignment of length 50, and so on. For our simulated datasets, where all sites evolved identically and independently, this simple method used to shorten genes is equivalent to selecting any subset of sites of a given length.

### Gene tree estimation

We estimated a gene tree on each shortened alignment with FastTree-2 version 2.1.8 [51], using the following command:

~~~
FastTreeMP -gtr -nt -gamma [input alignment]
~~~

We initially attempted to compute gene trees using RAxML; however, RAxML was unable to analyze many of the short sequence alignments we generated, because these lacked one or more nucleotides. Hence, we used FastTree-2, which does not have this problem. In addition, FastTree-2 is faster than RAxML, and prior studies have shown that trees computed using FastTree-2 are generally as accurate as trees computed using RAxML [38].

### Species tree estimation methods

*SVDquartets+PAUP\**

To run PAUP*’s version of SVDquartets, we used the following command within the command-line version of PAUP* 4.0a144 for Windows.

~~~
SVDQuartets nthreads=4 evalQuartets=all seed=5000;
~~~

For the mammalian biological dataset, we used PAUP* 4.0a142 for UNIX/LINUX with same command to run SVDquartets+PAUP*, but instead of using 5000 for the seed value, we used the default seed number, 0.

#### NJst

To run NJst, we used phybase version 1.4 [52] and custom scripts, available in our github repository.

#### ASTRAL-2

We used ASTRAL-2 version 4.7.6 [53], with the following command:

~~~
java -jar astral.4.7.6.jar -i [input gene trees] -o [output tree]
~~~

#### Concatenated analysis with RAxML

We ran an unpartitioned maximum likelihood (ML) analysis with RAxML, using the following command:

~~~
raxmlHPC-SSE3 -m GTRGAMMA -s [input alignment] -N 10 -p RANDOM -n [outputname]
~~~

Here, -`N` 10 indicates the number of starting trees for RAxML, and `RANDOM` is the random seed number, which we varied for each of the 10 runs.

In an unpartitioned ML analysis, all the sites in the aggregated superalignment are assumed to have evolved down the same model tree. Unpartitioned ML is a common approach to tree estimation, but is not as rigorous as a fully partitioned ML analysis, and has different theoretical properties in the presence of ILS [4].

The choice of unpartitioned analysis instead of fully partitioned analysis was necessary because RAxML cannot run if any of its parts (within a partitioned alignment) fails to have all four nucleotides present. While this constraint was not an issue for our longer gene sequence alignments, it caused substantial problems for more than half of the shortest gene sequence alignments.

### Evaluation Metric

All the estimated species trees returned by the analyses are bifurcating (i.e., all internal nodes have degree three). Hence, we report the Robinson-Foulds (RF) [39] error rate, which is equal to the missing branch rate for bifurcating trees. The script for this calculation is provided in our supplementary online materials.

#### Availability of supporting data

All datasets and supporting online materials are available at goo.gl/EgkWRk. A github repository containing all the source code used in our experiments can be found at https://github.com/j-chou/SVDquartets.git

#### Abbreviations

CA-ML: concatenated analysis using maximum likelihood
AD: average distance between true gene trees and true species tree
RF: Robinson-Foulds
GTR: Generalized Time Reversible ILS: Incomplete lineage sorting
MCMC: Markov Chain Monte Carlo

#### Authors’ contributions

JC computed gene trees, species trees using CA-ML, NJst, and ASTRAL-2 on the simulated datasets, ran SVDquartets+PAUP* on the biological dataset, analyzed data, made figures, wrote the first draft, and helped with the final draft. AG did some preliminary analyses on the simulated datasets, analyzed data, and helped write the first draft. SY wrote Python scripts, analyzed the data, and helped write the first draft. RD helped design the study, analyzed the data, and set up the github site. MN computed SVDquartets+PAUP* trees on all the simulated datasets, performed statistical tests, analyzed data, and made figures. SM simulated the data and helped with writing the first draft. TW conceived of the study, helped design the study, analyzed the data, and wrote the final draft.

## Acknowledgments

JC was supported by the Mathematics Department at the University of Illinois at Urbana-Champaign. RD was supported by NSF grant DMS-1401591. AG and SY were supported by the Computer Science Department at the University of Illinois at Urbana-Champaign. SM was supported by a graduate fellowship from the Howard Hughes Medical Institute (HHMI). MN and TW were supported by the National Science Foundation grant DBI-1461364.

